# Multiview tiling light sheet microscopy for 3D high resolution live imaging

**DOI:** 10.1101/2021.04.18.440322

**Authors:** Mostafa Aakhte, H.-Arno J. Müller

**Affiliations:** Developmental Genetics Group, Institute of Biology, University of Kassel, Heinrich-Plett Str. 40, 34132 Kassel. Germany

**Keywords:** microscopy, light sheet microscopy, selective plane illumination microscopy, Drosophila, early development, gastrulation

## Abstract

Light sheet or selective plane illumination microscopy (SPIM) is ideally suited for *in toto* imaging of living specimens at high temporal-spatial resolution. In SPIM, the light scattering that occurs during imaging of opaque specimens brings about limitations in terms of resolution and the imaging field of view. To ameliorate this shortcoming, the illumination beam can be engineered into a highly confined light sheet over a large field of view and multi-view imaging can be performed by applying multiple lenses combined with mechanical rotation of the sample. Here, we present a Multiview tiling SPIM (MT-SPIM) that combines the Multi-view SPIM (M-SPIM) with a confined, multi-tiled light sheet. The MT-SPIM provides high-resolution, robust and rotation-free imaging of living specimens. We applied the MT-SPIM to image nuclei and Myosin II from the cellular to subcellular spatial scale in early *Drosophila* embryogenesis. We show that the MT-SPIM improves the axial-resolution relative to the conventional M-SPIM by a factor of two. We further demonstrate that this axial resolution enhancement improves the automated segmentation of Myosin II distribution and of nuclear volumes and shapes.

## Introduction

The early development of multicellular animals features dramatic tissue movements resulting in the spatial organization of the body plan. An important mechanism driving these dynamic cell and tissue movements are force-producing molecular interactions and their mechanobiological responses [1]. In an ideal scenario, the molecular dynamics underlying cell and tissue movements should be investigated in the context of the entire embryo, because all cells are mechanically coupled to some extent. To achieve this goal, robust and fast microscopy methods are required to dynamically image entire developmental processes. The prevailing method for most of these studies in embryogenesis has been confocal-based fluorescence microscopy, because of its high spatial resolution in three-dimensional multi-channel imaging. These applications have provided important information on the biophysics of actomyosin dependent dynamics related to tissue tension and other mechanical feedback within a tissue [2]. However, confocal microscopy of whole embryos is limited by a rather small field of view and the relatively high level of irradiation during the scanning process bearing the risk of phototoxicity. Therefore, it still remains a challenge to image all morphogenetic movements in an embryonic phase like gastrulation simultaneously and at a subcellular level of optical resolution.

One method that is specifically well suited to address the challenges in dynamic imaging of whole embryos is selective plane illumination microscopy (SPIM). SPIM bypasses some of the limitations of the conventional epifluorescence confocal microscopy [3]. The major advancement of the SPIM in comparison to epifluorescence microscopy lies in its ability of imaging with minimal out-of-focus light and low phototoxicity. These improved imaging parameters are achieved by applying a thin sheet of light to illuminate the sample. The simplest platform of the SPIM consists of two separate optical arms: The illumination arm and the detection arm, which are arranged at an angle of 90 degrees to each other [3, 4]. The illumination arm produces a sheet of light that illuminates only one plane within the sample, a method also known as optical sectioning. The performance of the optical sectioning of a given light sheet is defined by its thickness and its effective range. The effective range of the light sheet determines the imaging field of view (FOV) and these parameters have an inverse relationship with each other [5]: a thick light sheet has a long effective range, but provides poor optical sectioning; a thinner light sheet provides better optical sectioning, but results in a smaller FOV. The ideal set-up should take these parameters into account and will apply a thin light sheet over a large FOV (**Supp. Mat. Fig. 1**).

**Figure 1.**
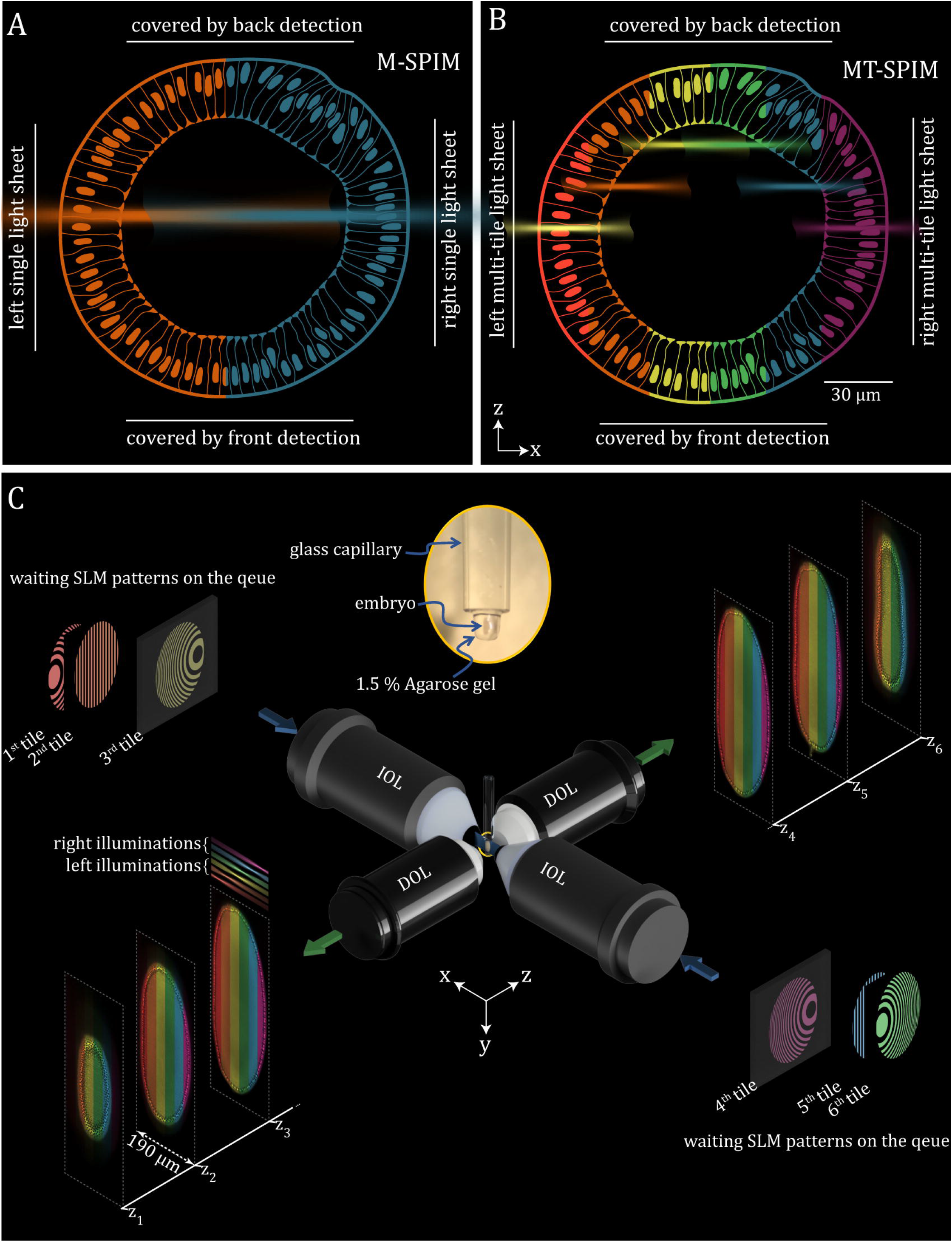
Principle of Multiview tiling selective plane illumination microscopy. **(A)** Concept of the conventional Multiview selective plane illumination microscopy (M-SPIM) from the axial view (xz plane). The sample is illuminated with two homogeneous and thick light sheets from opposite directions, which are color-coded with blue and orange. **(B)** Design of Multiview tiling selective plane illumination microscopy (MT-SPIM) with six tiles of illumination in which each tile covers 16.6% of the volume. **(C)** The workflow of the MT-SPIM which consists of four objective lenses; two illumination and two detection lenses. The Illumination arms includes a spatial light modulator with the combination of two Galvo mirrors (GMy and GMz) to tile, generate and swipe the light sheet in three dimensions through the sample, which is dipped into the water between four objective lenses.

The SPIM’s performance is not only limited by the properties of the light sheet itself, but also by the optical properties of a biological sample causing light scattering. Using wavelengths typically used for fluorescent imaging, the light sheet does not penetrate more than several tens of micrometers into the living biologicals sample which greatly affects the quality imaging [6]. This is particularly true for fairly large and opaque specimens, in which the light sheet is heavily diffracted over the imaging FOV, resulting in only a small fraction of the returning well-resolved information [3]. Multiview imaging represents one efficient approach to compensate for poor optical sectioning and sample transparency [4, 7]. The specimen is rotated mechanically in order to illuminate and image the sample from different angles. The speed of the mechanical rotation is limited to avoid a drift of the sample resulting in a blurred or de-positioned image. As a consequence, the slow speed rotation creates a time delay between the different views during 3D image capturing. Moreover, if the mounting of the sample is not centered to the axis of rotation, it creates an additional lateral or axial shift during imaging for each rotated view. These limitations can be addressed by re-arranging the basic Multiview imaging to a system with four optical arms consisting of two illumination arms and two detection arms [7, 8]. The illumination arms create two light sheets which excite the sample from opposite sides and each light sheet covers half of the sample. The emitted light from each FOV is collected by two objective lenses which are arranged orthogonally to the light sheets. Compared to the basic Multiview imaging setup, the Multiview SPIM with four objective lenses is at least four times faster and it provides less blurring effects during imaging, thus improving reconstruction and quantification of the imaging data.

Another key factor related to the FOV in Multiview SPIM is the axial resolution of the microscope. Similar to the basic SPIM configuration, the axial resolution of the Multiview SPIM can also be defined by convolution of the light sheet and the point spread function of the detection lens. Two principal approaches can improve the axial resolution of a Multiview SPIM: sample rotation and light sheet engineering. As described above, the rotation allows for covering the entire 3D FOV within the SPIM setup. In the case of the resolution enhancement with a rotation approach, the four-lenses Multiview SPIM also can be combined with the sample rotation technique such that each quarter of the sample is imaged at least two times from two orthogonal views. This sequential imaging of the sample causes a time delay between two views of each part that may result in a misaligned image fusion when live imaging of subcellular activity is concerned [8]. Previous studies show that these limitations can be addressed by a complicated Multiview SPIM arrangement in a way that each optical arm functions as illumination and detection at the same time [9]. This helps to increase the accuracy of the imaging process, however, it requires an expensive and large optical setup with four cameras and eight scanning mirrors.

The lack of axial resolution of a light sheet over a large FOV can also be compensated for using the so-called tiling technique, which involves manually moving the sample through the thin light sheet or sweeping multiple thin light sheets inside a large sample [10–13]. The light sheet tiling method has the advantage over moving the sample, because it is faster and contact-free. The tiled light sheet can be generated by rapidly adjusting the wave-front of the excitation beam using several beam shaping approaches: an electrically tunable lens [12], a tunable acoustic index gradient lens [10,13], mechanical remote focusing by a piezo electric element [11] or a fast spatial light modulator (SLM) [14]. Among these methods, the SLM is the most practical beam shaper, because it offers a versatile application for beam shaping as well as beam aberration correction and beam aperture controlling simultaneously with the tiling technique [15]. All of these approaches were successfully applied to the basic configuration of the SPIM with two or three objective lenses [10, 11, 13, 14, 16–21] to imaging a small transparent specimen or cleared fixed sample, but are not optimized for high resolution whole functional imaging of an opaque live sample.

In the present work, we combined the simultaneous M-SPIM with the tiling light sheet technique to create a novel robust SPIM for high resolution live imaging. In our technique, a large and opaque specimen like the *Drosophila* embryo can be imaged from four different views with high spatio-temporal resolution without any rotation. First, we demonstrate the concept of MT-SPIM based on the implication of a fast SLM to tile the scanning light sheet. The broad versatility of MT-SPIM is demonstrated by live imaging of MyoII-GFP in early *Drosophila* embryos. We compared the axial resolution of the recorded data from the tiling MT-SPIM mode with data obtained by the conventional, non-tiling M-SPIM mode. The MT-SPIM technique yields images with a high spatial resolution in 3D over time, which can be employed to reconstruct the natural shape of each cell in a fairly large multicellular organism. We show that 3D imaging with MT-SPIM of the cell nuclei greatly improves automatic segmentation and analysis of nuclei number and volume. We analyzed the imaging data to correlate the dynamics of subcellular MyoII distribution with supracellular MyoII structures within the context of the entire embryo. This analysis revealed the temporal sequence of dynamic MyoII-GFP localization in distinct morphogenetic movements during gastrulation in the *Drosophila* embryo.

## Results and Discussion

### Design of the MT-SPIM microscope set-up

The conventional multi-view light sheet microscopy with four objective lenses is based on simultaneous or sequential light sheets to illuminate the sample from two sides [7, 8] (**Fig. 1A**). The dimensions of these two light sheets are usually determined by the size of the sample, thus that ideally each light sheet should cover half of the sample. The standard and commonly used parameter to define the light sheet’s FOV is twice the Rayleigh length, which is the distance from the smallest waist of the light sheet to the point where it becomes 1.4 times wider than its center. Therefore, an inherent feature of the light sheet is a non-uniform axial resolution that results in differences of the imaging quality over the FOV (**Suppl. Fig. S1)**. For instance, a light sheet based on a Gaussian beam needs to cover a 90 μm FOV, when imaging half of the diameter of a *Drosophila* embryo. Accordingly, the waist of this light sheet is about ∼2.7 μm within the central area, but it becomes ∼3.8 μm at the edges causing a loss in axial resolution. To achieve higher levels of resolution, the waist of the light sheet should be even thinner, which on the other hand aggravates the compromised optical sectioning at the edges (**Suppl. Fig. S1A,B)**. In order to overcome this inhomogeneity in optical sectioning, we combined the M-SPIM with a beam tiling method based on a SLM (**Fig. 1B**).

The optical design of our MT-SPIM set-up is based on a classic arrangement with four objective lenses, where the sample can be dipped into water in the space between the lenses within a sample chamber (**Fig. 1C**). The sample and illumination objective lenses are in a fixed position and are not moving, which protects the imaging from adverse effects through unwanted movement. The optical illumination paths include a fast SLM and two scanning mirrors: a Y-Galvo mirror and a Z-Galvo mirror. The Y-Galvo mirror scans the illumination beam along the height of the imaging FOV (Y-axis) to generate the light sheet, and the Z-Galvo mirror scans the light sheet though the sample for volume imaging. At the same time, the SLM modulates the illumination beam several times per each imaging plane by applying virtual lenses to tile the light sheet in a remote fashion through the X-axis (**Fig. 1C; Suppl. Fig. S2**). Each illuminated tile of the sample is recorded independently through rapidly changing the SLM’s pattern which has a 41 μs time gap between two tiled light sheets. This enables us to rapidly swipe the light sheets over the FOV for near simultaneous illumination. The combination of this illumination approach with Multiview-SPIM enables us to apply a thin light sheet over the entire *Drosophila* embryo. The temporal resolution of the MT-SPIM supports the time-lapse recording of most morphogenetic movements in 3D.

### Enhancing resolution by MT-SPIM imaging

The MT-SPIM setup provides improved dynamic imaging properties by combining a high spatial and temporal resolution. To investigate the properties of the MT-SPIM microscope, we performed a set of experiments by imaging 2 hours-old, living *Drosophila* embryos expressing Sqh::eGFP (regulatory light chain of non-muscle Myosin II [MyoII] fused to eGFP). The early *Drosophila* embryo is a large, diffractive sample that requires a 190×190×500 μm^3^ volume FOV for imaging. In order to compare the performance of the MT-SPIM in comparison to the conventional M-SPIM mode, we applied two distinct types of light sheets during imaging simultaneously. Both light sheets were based on a scanning Gaussian beam: The conventional light sheet waist was uniform along the FOV (around 90 μm) and its full width at half maximum (FWHM) was about 3.2 μm. For the tiling light sheet, we used a Gaussian light sheet with a FWHM on its waist of about 2.2 μm which has a 65 μm effective length (two times of Rayleigh range) where each single tiling light sheet is non-uniform and its Rayleigh range covers less than half of the embryo (**Suppl. Fig. S3**). This means that its waist in the center (2.2 μm) is smaller than to the edges (3 μm), which makes better light confinement and axial resolution in the center of FOV than to the edges. To compensate the resolution and contrast on the FOV’s edges, we have tiled the light sheet several times per illumination arm. For instance, to deliver a homogeneous light sheet over the embryo, we have tiled the light sheet three times for each illumination arm (**Suppl. Fig. S4**). The center of each tile has 30 μm displacement (one Rayleigh range) from the next tile, which creates a sufficient amount of overlap between the tiled light sheets to generate a homogenously illuminated area in the MT-SPIM configuration (**Suppl. Fig. S5)**.

One advantage of the SLM-based Multiview design is to rapidly switch between tiling and non-tiling modes. Thus, a sample can be imaged with different beam engineering techniques simultaneously, which allows a side-by-side comparison of data with M-SPIM and MT-SPIM from the same embryo (**Fig. 2A**). To examine the microscope’s resolution in the axial direction (Z-axis), the 3D recorded images were reconstructed as maximum intensity projections in the corresponding axial direction (**Fig. 2B**). The side-by-side comparison of various cellular structures of the Sqh::eGFP embryo in different regions of interest (ROI) illustrates the improved resolution of the dynamic MyoII-GFP localization (**Fig. 2C**). The Fourier analyses confirm that the frequency components of the MT-SPIM image are more expanded in the Z-axis compared to the M-SPIM image; we calculated a twofold improvement of the Z-resolution in the tiling mode when compared to the M-SPIM mode **(Fig. 2D, MOVIE 1-2)**. The improvement in axial-resolution is critical for the dynamic visualization of subcellular components in three dimensions. For instance, transversal cross-sections of embryos can reveal morphogenetic events in the internal of the *Drosophila* embryo to analyze the localization of MyoII-GFP from dorsal, ventral and lateral views of the embryo at the same time (**Fig. 2E**). The 3D-rendered data (**Fig. 2**) show that the MT-SPIM mode improves the reconstruction accuracy of subcellular details. The subcellular MyoII network is visible at higher resolution not only at the embryo surface but also when the actomyosin network moves into the interior of the embryo during cell formation (**Fig. 2E**). The MyoII localization around each cell during this rapidly advancing dynamic process can be recovered more precisely by MT-SPIM (**Fig. 2E)**.

**Figure 2.**
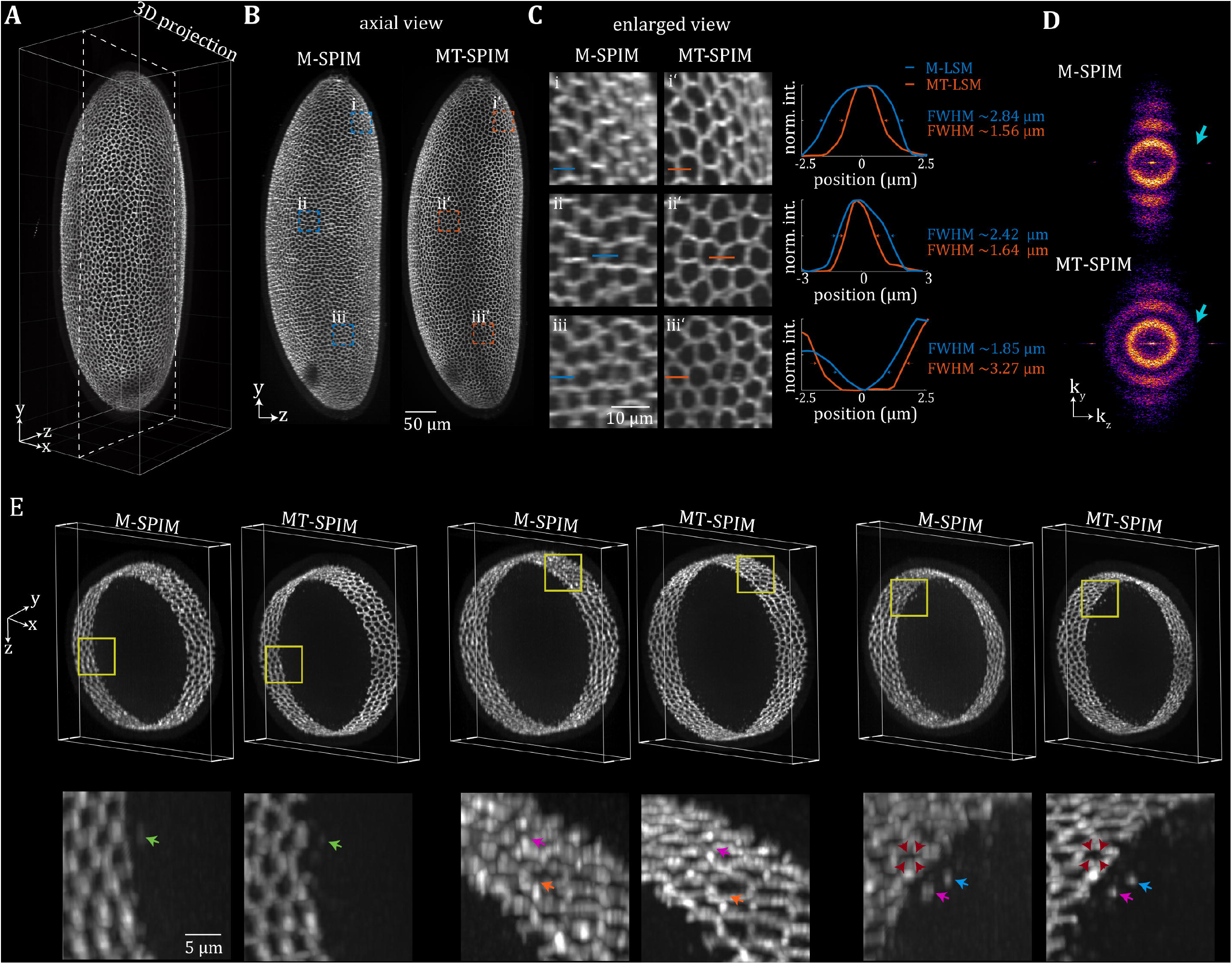
Improvement of three-dimensional image resolution with MT-SPIM microscopy. **(A)** 3D rendered image of the whole *Drosophila* embryo expressing Sqh-GFP (non-muscle Myosin II tagged with GFP). **(B)** Axial view (YZ plane) maximum intensity projection of the embryo from the conventional M-SPIM and MT-SPIM. **(C)** Side by side comparison of the of enlarged three regions of interest from images in **(B). (D)** Calculated frequency domain of the images in **(B)**, which illustrates the higher spatial frequency in z-direction from the MT-SPIM imaging compared to the conventional M-SPIM. **(E)** This panels shows the 3D rendering of the *Drosophila* embryo from its small cross-section from three different 3D ROIs. The arrows highlight the difference between MT-SPIM and M-SPIM to reconstruct the MyoII accumulation and its natural shape.

### Live imaging of early morphogenesis stage in Drosophila embryos using MT-SPIM

The MT-SPIM is designed for high resolution 3D imaging of fairly large specimens over time. The lateral and axial resolution of the MT-SPIM is measured at 800 ± 0.04 μm and 1.4 ± 0.09 μm with deconvolution, respectively. The imaging speed for the whole embryo volume is 40 seconds to capture 1300 image frames where the exposure time is set to 20 ms. To explore the full potential of our MT-SPIM method for live imaging we investigated the dynamic MyoII-GFP distribution during early development. As a proof of principle, we acquired 3D image sequences of 2 hours-old live *Drosophila* embryos expressing Sqh::eGFP every 90 seconds (80 seconds for simultaneous imaging of tiling and conventional modes and 10 sec interval time) for 90 minutes during cellularization and the beginning of gastrulation (**Fig. 3**). During this early developmental stage, actomyosin forms a polygonal network over the entire embryo, which moves towards the center of the embryo [22]. During this inward movement, the actomyosin network changes its configuration starting with a priming phase to a hexagonal phase and finally ending in a ring phase where individual actomyosin rings contract [23].

**Figure 3.**
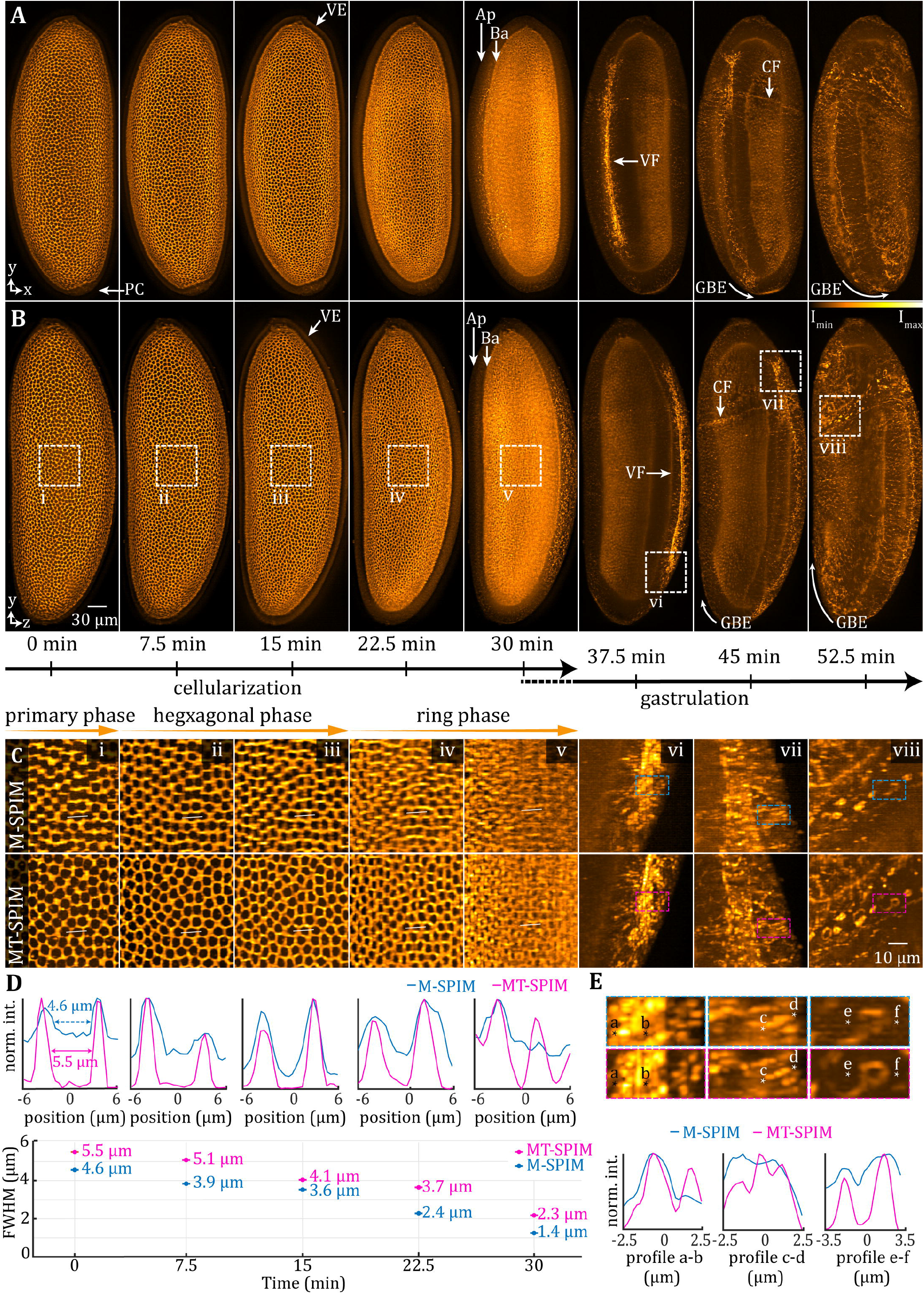
Side by side comparison of axial resolution performance from MT-SPIM and conventional M-SPIM mode in live 3D imaging. **(A)** Maximum intensity projection of images (xy plane) recording MyoII-GFP during the early development of the *Drosophila* embryo in the MT-SPIM mode. A temporal series is shown (from left to right) representing embryonic stages from cellularization to the onset of the germband extension. **(B)** Reconstructed axial views of the images in (**A**). The arrows highlight different regions of the embryo such as PC: pole cells, VE: vitelline envelope, Ap: apical aspect of the cells, Bp: Basel aspect of the cells, VF: ventral furrow formation, CF: cephalic furrow formation, GBE: germband extension. **(C)** The boxed areas represent magnified regions of interests (ROIs) (i-viii) from (**B**) to compare the performance of MT-SPIM and M-SPIM when reconstructing the axial resolution of the live imaging. **(D)** Comparison of normalized intensity profiles from the images (**C**). The pink and blue lines show the profile intensity of the selected line from MT-SPIM and M-SPIM (white lines in (**C**), and the space between two peaks show the distance of two opposite cell contacts. The average of the measured distance indicates a larger separation (about 1 μm) between two opposite cell contacts from the MT-SPIM method compare to the M-SPIM method. **(E)** Enlarged views of (**Cvi-viii**), which show the capability of the MT-SPIM compare to conventional M-SPIM to reconstruct the nmMyoII-GFP accumulation in ventral furrow formation and contractile ring formation during cell division.

The lateral (XY plane) and axial views (YZ plane) of the sample were reconstructed for each time point by fusing the tiles using content-based image fusion (**Fig. 3A-B**). The improved optical sectioning capability and the removal of out-of-focus signals are apparent in the MT-SPIM images in the lateral and in the axial direction (**Fig. 3A,B; MOVIE 3)**. A side-by-side comparison of the reconstructed data indicated artifacts in distribution of MyoII-GFP in the M-SPIM data during the different phases of cell formation; MyoII-GFP is lacking at several anterior-posterior cell boundaries suggesting a partially planar polarized distribution at the onset of cellularization (**Fig. 3Ci-v**). When comparing the average distance between two opposite bi-cellular contacts at an identical position, it became apparent that the MT-SPIM resolution is improved as the distance appears 1 μm shorter when compared to M-SPIM (**Fig.3Di-v)**. This difference can become even more extreme when the network enters the ring phase, where the individual rings cannot be resolved in M-SPIM mode, but can be imaged as discrete structures in MT-SPIM imaging (**Fig. 3Cv,D**). The MT-SPIM microscope also greatly improved the resolution when imaging various subcellular and supracellular MyoII-GFP structures that form during gastrulation movements (**Fig. 3E**). We conclude that the increased axial resolution of the MT-SPIM is critical to provide an improved reconstruction of subcellular MyoII-GFP structures within the context of the entire development of the early *Drosophila* embryo.

### 4D dynamic imaging of MyoII-GFP distribution during selected morphogenetic movements in the *Drosophila* gastrula

The MT-SPIM imaging of provides an excellent method to study the dynamic changes in the distribution of GFP-tagged proteins in the context of early *Drosophila* embryos. A whole range of different morphogenetic movements have been well studied during gastrulation and the actomyosin cytoskeleton was shown to be involved in most of these events [24]. The simultaneous imaging of MyoII-GFP in the embryo at the mesoscale has provided important information about the relationship of mechanical forces in instructing cell flows during gastrulation movements in *Drosophila* [25]. While individual subcellular and supracellular MyoII structures have been detailed by high-resolution confocal microscopy, these structures have not been imaged in the context of the entire early embryo, including internal tissues. We therefore applied the MT-SPIM to dynamically image the distribution of MyoII-GFP and quantified fluorescence intensities in selected events during gastrulation (**Fig. 4**).

**Figure 4.**
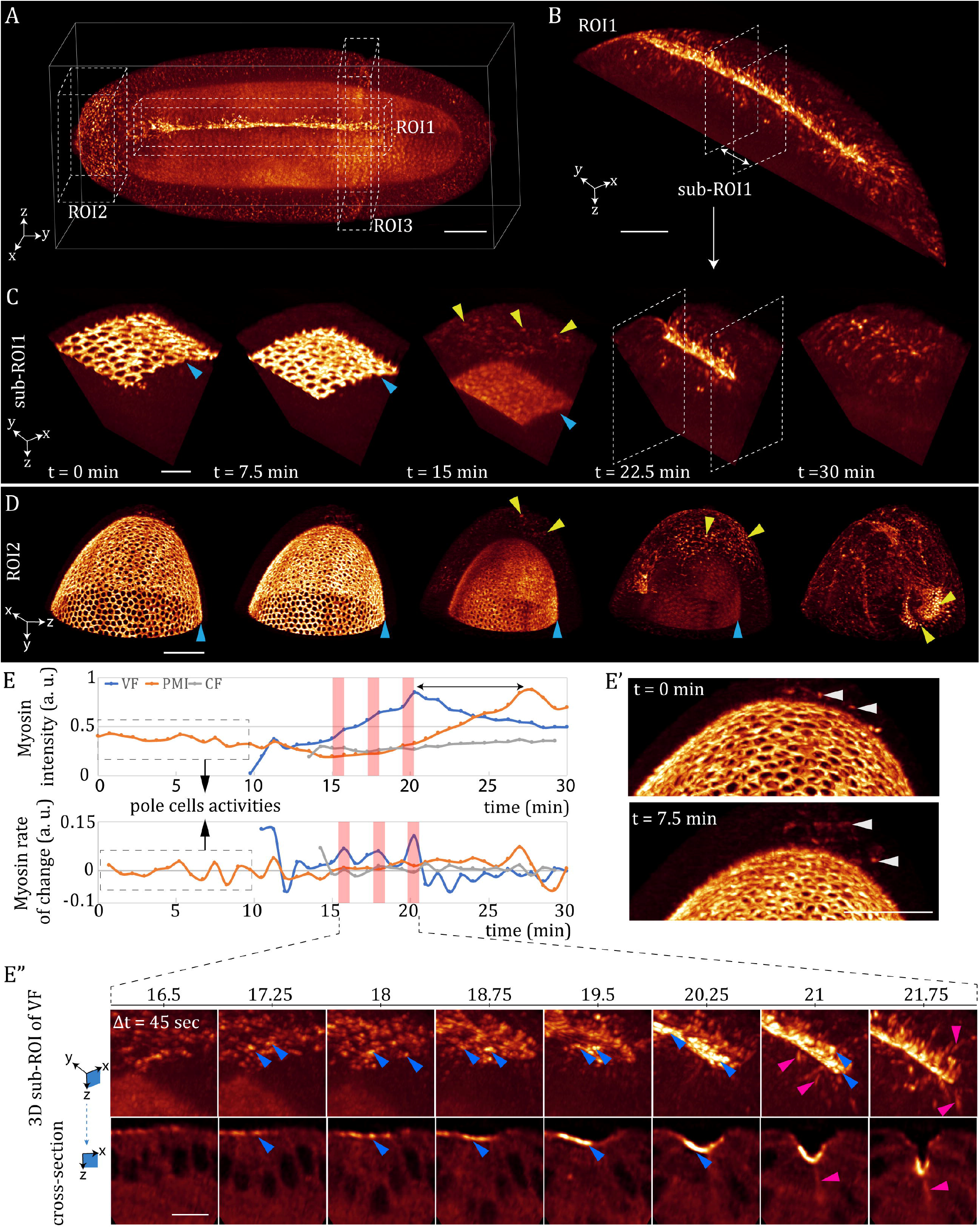
The capability of the MT-SPIM for simultaneous 4D quantitative data analysis of subcellar MyoII-GFP distribution during *Drosophila* gastrulation. **(A)** 3D rendering of the whole *Drosophila* embryo with MyoII-GFP recorded with MT-SPIM using six tiled light sheets. Three regions of interest are selected for simultaneous investigation, ROI1: ventral furrow, ROI2: posterior midgut, ROI3: Cephalic furrow. **(B)** MyoII dynamics on the ventral side during furrow formation visualized by a cropped 3D region of interest of (**A**). The bright line, which is elongated from anterior to posterior, shows the MyoII-GFP accumulation from apically constricting cells in the centre of the ventral furrow formation. **(C)** 3D region of interest from the middle part of the (**B**) during embryo development. This sub-region of interest illustrates the embryo development from cell formation and early gastrulation. The basal domains of the cells are indicated by the blue arrow indicating the MyoII movement from apical to basal. The yellow arrows indicate the MyoII activity in the apical domain of the cells coinciding with end of the cell formation. **(D)** 3D region of interest from the posterior side of the embryo during cellularization and early gastrulation. The blue arrows show the basal part of the cells during cellularization, and the green arrows indicate the myosin activity in the apical part of cells in the posterior. **(E)** The quantification of the myosin intensity in 4D for apical side of the cells for the three regions of interests (ROI1, ROI2 and ROI3) of the embryo that has shown in (A-D). The first graph shows the mean value of the MyoII-GFP fluorescence intensity at the apical domain of the cells for each ROIs (the basal part is excluded), and the second graph describes the MyoII-GFP rate of change along the time. The first 10 minutes of the recorded data is indicated with a dashed rectangle in the graph illustrates. This part shows that the only signal that can be extracted at this time is related to the pole cell activities in the posterior side of the embryo (see white arrows in **E’**). The intensity in the apical part of the ventral side is increased with a specific oscillation frequency. The pink areas in the graphs show the myosin activities’ peak during the early gastrulation, which is indicated in detail in the subset figures (**E”**). (**E”**) the first row of this figure shows the 3D rendering of the myosin activities in a small part of the ventral side during the ventral furrow formation. The bright spots which are indicated with the blue arrows are the medial myosin on the apical side. The pink arrows also show some of the myosin-like cable structures necessary for apico-basal traction for ventral furrow invagination. Scale bars are 50 μm for A, B, D, E’ and 10 μm for the C and E’’.

During cellularization the polygonal actomyosin network moves from the periphery of the embryo towards the centre. Towards the end of cell formation, the maximum depth of the network varies along the periphery of the embryo; we measured a depth of about 42 ± 0.1 µm in the middle section and 24 ± 0.25 µm at the poles (**Fig. 4C,D)**. We quantified the MyoII-GFP intensities for three well described morphogenetic movements during gastrulation: ventral furrow formation, posterior midgut formation and cephalic furrow formation (**Fig. 4B**). During ventral furrow formation, MyoII-GFP intensities at the apical domains of the presumptive mesoderm cells arise at about 10 min after start of the cellularization and then gradually increase during early gastrulation. Synchronously, the MyoII-GFP intensities at the cephalic furrow begin to increase, but to a much slower extent compared to the ventral domain. At the posterior pole the first fluctuations in MyoII-GFP intensities were observed before the onset of gastrulation, which were caused by the mitotic activities of the pole cells (**Fig. 4E, E’**). A strong MyoII-GFP intensity increase is observed in the posterior midgut invagination with 7.5 minutes delay after the highest peak of MyoII-GFP in the presumptive ventral furrow. These data show that the first changes in MyoII-GFP occur in the ventral domain followed by the cephalic furrow and last in the domain of posterior midgut formation.

The data also allowed us to calculate the rate changes of MyoII-GFP intensities for the individual morphogenetic movements (**Fig. 4E)**. Such analyses may reveal whether the MyoII changes occur by similar principles or whether there are potential differences in the mechanisms of MyoII accumulation. We found that MyoII-GFP rate changes followed a different pattern between the different morphogenetic events. In the cephalic furrow area the increase in MyoII-GFP levels occurred without any notable rate changes. In contrast, the increase in the ventral furrow showed alternating phases of high and low rates of intensity changes. The posterior midgut exhibited also alternating rate changes, however with a longer amplitude than the ventral furrow. These results indicate that rate changes underlying the increase in MyoII-GFP levels in different morphogenetic movements are different from each other.

The differences in the rate changes of MyoII-GFP intensities suggest different mechanisms or differences in the MyoII associated mechanics in these movements. These differences may be due to the distinct geometries of the position where the fluctuations occur within the embryo. In case of the ventral domain, it was striking that the rate changes in the ventral furrow formation reflect in timing and spatially the reported pulsed contractions that drive this process [26]. While these pulsatile behaviors have been well described on subcellular levels, here we provide evidence that the changes in the rate of MyoII increase in the entire ventral furrow domain displays fluctuations that occur within the very similar time frames as shown for small groups of cells.

We conclude that MT-SPIM imaging is able to image dynamic subcellular events within the context of the entire embryo. Furthermore, the fact that rate changes exhibit an oscillatory behavior indicate that they are not random over the domain of the ventral furrow and the posterior midgut. The frequencies of the rate changes were comparable to the MyoII-GFP rate changes measured by confocal microscopy. This supports the findings that the contractions are coordinated over the entire range of the ventral furrow in this phase of ventral furrow formation [27].

### Live quantitative imaging of nuclei forming during cellularization in the *Drosophila* embryo by MT-SPIM

We next examined whether the performance of the MT-SPIM is sufficient for the imaging and segmentation of the nuclei in the entire embryo. To investigate the effect of the improved resolution on the reconstruction of the nuclei, we imaged a 2h old *Drosophila* embryo expressing His2Av-GFP by MT-SPIM and M-SPIM simultaneously with six tiled light sheets. His2Av-GFP is also found within more central areas of the embryo, where it is present in lipid droplets that store histones for early embryogenesis (**Fig. 5B; Movie 6**) [28]. The Multiview image fusion let us reconstruct the embryo’s nuclei from any arbitrary plane, such as long and small cross-sections (**Fig. 5A-C; Suppl. Movie 6**). However, the nuclear shape and the border between them were obscured in M-SPIM data along the axial direction. In contrast to the M-SPIM, the MT-SPIM is able to resolve individual nuclei even from an axial view. Therefore, the quality of the reconstructed nuclei is completely dependent on the area selected for the analysis and for instance, the nucleus located in the axial imaging view cannot be distinguished easily as an individual nucleus using the M-SPIM method (**Fig. 5D**). To explore the effect of the improved resolution, the data recorded from either SPIM set-up were segmented based on the automatic thresholding algorithm. The segmentation was done for a region of interest (ROI) and the whole embryo over the cell formation (**Fig. 5E**). The results show that the two imaging modes predict different numbers of nuclei in the embryo. The number of the nuclei within the ROI is about five times higher in data based on the tiling mode compared to the conventional imaging mode. We also measured each nuclei volume for the ROI and whole embryo. The results show that the volume measurement is also depended on the quality of the segmented nuclei. The averaged nuclei volume by the conventional light sheet was measured between 135 ± 275 µm^3^and 176 ± 10 µm^3^over cellularization, compared to the measurements based on the MT-SPIM data that were between 122 ± 20 µm^3^and 160 ± 4 µm^3^. The lower standard deviation in the MT-SPIM data shows that more volume nuclei are clustered around the average size, while the M-SPIM data shows a larger error in measuring nuclei volume over cellularization, which is caused by the poor axial resolution. We conclude that the quality of the image reconstruction of M-SPIM *vs*. MT-SPIM data can differ to an extent that may directly affect the biological interpretation.

**Figure 5.**
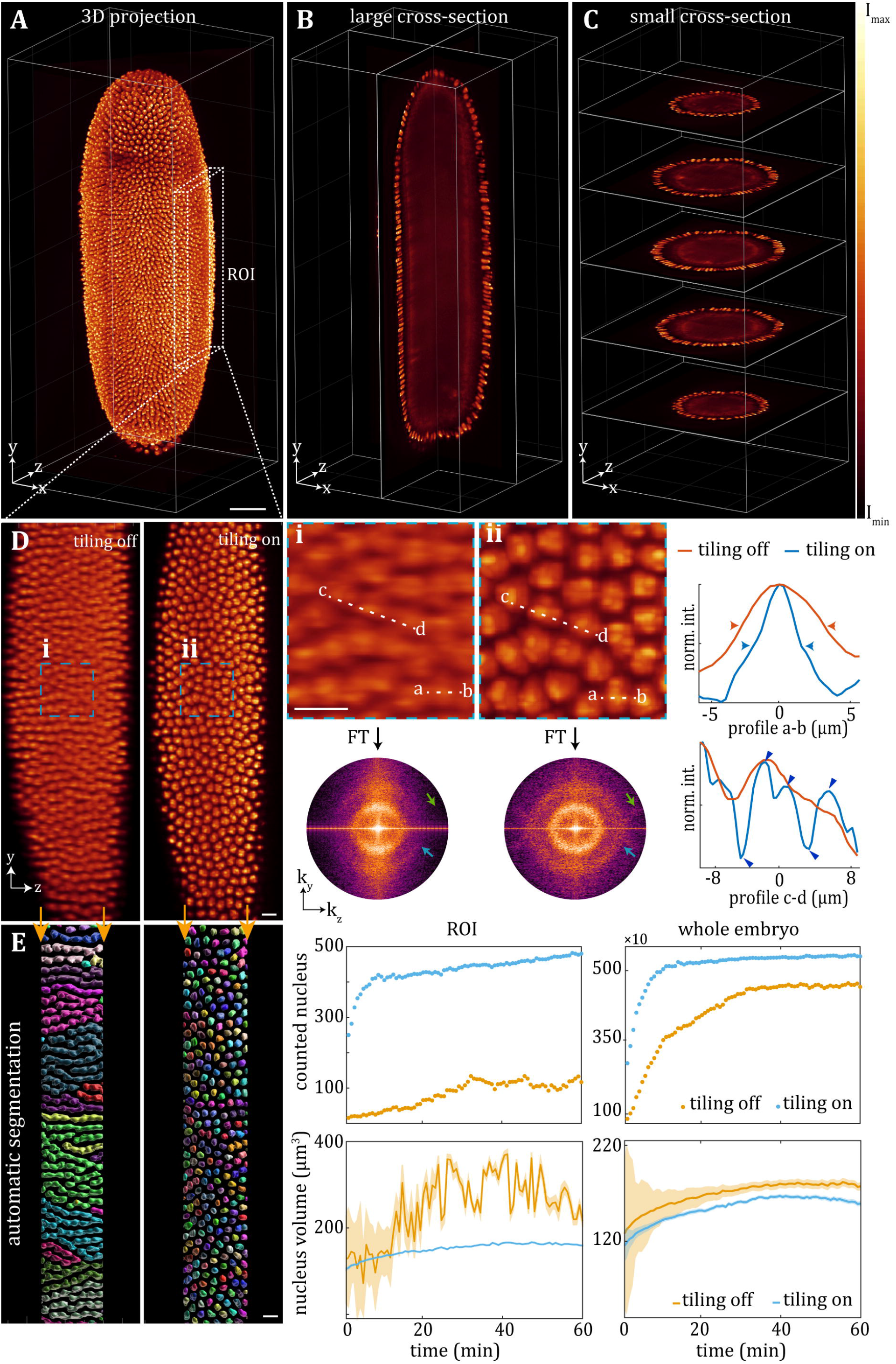
The performance of the MT-SPIM to 3D-imaging of cell nucleus. (**A**) 3D rendering of the whole *Drosophila* embryo with His2Av-GFP. The x-y axis indicates the lateral imaging plane, and the y-z and x-z planes show the axial imaging planes. (**B,C**) High quality 3D-rendered data enables us to study the different cross-sections from arbitrary views. In addition to the nuclei,His2Av-GFP can also be seen in lipid droplets beneath the cortical cytoplasm (**D**) A region of interest (ROI1) from (**A**) is selected for the first step of the resolution analysis over the axial view for both Multiview imaging modes, tiling and non-tiling. (**Di** and **Dii**) Higher magnification of the dashed rectangular regions are provided to show the differences between the reconstructed data from both imaging modes. The intensity analysis show when the tiling mode is turned on the size of each nuclei is reconstructed 1.5 times smaller compare to non-tiling light sheet which shows the capability of the tiling mode to distinguish the adjacent nucleus. In addition, the Fourier analysis verifies the resolution enhancement over the axial direction when the tiling mode is applied. (**E**) 3D-automatic segmentation for the central part of the images in (**D**) are provided to illustrate the effect of the resolution on automatic segmentation. (**F**) The number of the nucleus and the volume of each nuclei is also calculated based on the automatic segmentation for ROI1 and whole embryo over cellularization. Scale bars are 50 μm for **A-B** and 10 μm for the **D-E**.

## Conclusion

MT-SPIM provides a rotation-free and compact optical setup for simultaneous subcellular and cellular imaging of a multicellular and opaque live organism. This technique aids in determining the properties of a subcellular organelle within a sample’s region of interest and its dynamic changes over time in various regions of the sample. The time resolution of the imaging represents a crucial parameter in light sheet imaging and may be compromised by the beam tiling as a function of the number of tiles applied, the exposure time, and the inter-tile delays. Hence, we have optimized this system by using a fast SLM which allows illumination of the sample in a near-simultaneous way for each frame. The imaging speed of the MT-SPIM can be adjusted by the recording time of each tile associated with the SLM and the camera exposure time. Weak fluorescent signals will require a longer exposure time, which can be accomplished by repeating the light sheet in each tile modulated by the SLM. These adjustments can easily be achieved to detect sufficient signals in diverse applications in order to resolve time and spatial resolution issues. With that given time and spatial resolution, we have shown that we are able to record protein dynamics such as of Myosin II on a subcellular scale within the whole embryo without the need of rotating the sample. Also, we have shown that the MT-SPIM results can be combined with automatic segmentation to predict the volume and number of the nuclei in early embryos.

## Material and methods

### Multiview tiling-SPIM microscope configuration

In the proposed Multiview tiling-SPIM, a diode laser with 488nm wavelength operates as an illumination source to excite the GFP based sample. The laser beam’s output diameter of about 1 mm is insufficient to interact with a spatial light modulator (SLM) (SXGA-3DM-DEV Microdisplay device, Forth dimension displays) with a diameter of about 12 mm. As a result, the laser beam is expanded to a diameter of 12 mm using a two-lens telescope configuration (L1: focal length f = 25 mm, L2: f = 300 mm) to cover the entire region of the SLM for efficient beam shaping. The reflected light from the SLM is then passed through a lens (L3: f = 100 mm), resulting in a Fraunhofer diffraction pattern in its Fourier plane. The unwanted diffraction components caused by the inherent pixelated SLM structure are filtered by a pinhole. Following the pinhole, the filtered light is collected by another lens (L4: f = 75 mm) to create a conjugation of the pattern projected by the SLM. The light then hits a series of optical components, including a Z-Galvo mirror, two lenses, a Y-Galvo mirror, and a Fourier transforming lens. The Z-Galvo mirror (GVS412/M, Thorlabs) is used to reposition the illumination light through the sample for volume imaging, which is conjugated to the Y-Galvo mirror (GVS412/M, Thorlabs) by the pair lenses to create the light sheet. The primary light sheet is detectable in the focal length of the Fourier transforming lens (L6: f = 75 mm) positioned in front of the Y-Galvo mirror. The primary light sheet is divided by a knife edge mirror (MRAK25-G01, Thorlabs) to direct the light sheet into the right and left illumination arms in order to achieve two side illumination. The shaped primary light sheet is then collected by a lens (L7: f = 150 mm) and an objective lens (N10xW-PF, 0.3 NA, 3.5 mm WD, Nikon) assembly, which guides the light sheet to the chamber to excite a particular layer of the sample. The emitted light is then collected from two directions at the same time by two objective lenses (20xW, Water, 0.5 NA, UMPLFLN, Olympus). A quick piezo positioner is used to align the imaging objective lenses with the position of the light sheet (P-625.1CD, PI derived by E-709.CHG). The collected light is then imaged on a single sCMOS camera (Orca-flash 4.0 V2, Hamamatsu) with a knife edge mirror and a combination of two tube lenses (U-TLU-1-2, 200 mm, Olympus).

### Multi-tiled light sheet based on SLM

To enable the SLM to be used for tiling, it is encoded with several patterns that allow for the focal point of the light sheet to be shifted along the direction of the beam propagation. The SLM operates on the basis of a digital pattern referred to as a hologram, which is usually synthesized digitally. Specifically designed for tiling, a digital hologram is a pattern that behaves as a lens. The ability to encrypt a lens into a programmable SLM is appealing since the focal length of the lens can be varied at the frame rate of the SLM, allowing access to a rapidly tunable lens. To implement this method and create the tiling light sheet, we used the SSPIM toolbox to create digital holograms based on virtual lenses. [15].

### Microscope control and operating system

In order to high-speed imaging the Multiview tiling-SPIM is controlled by a fast programable gate array (FPGA) (PCIe-7851 Multifunction Reconfigurable I/O Device, National instrument) (**Supp. Mat. Fig. 4-5**). The microscope is controlled with a home built graphical user interface (GUI) based on LabView 64-bit which is running on a high-speed computer with 128 GB random-access memory (RAM), 18 central processing units (CPU) (3 GHz) and 8 TB solid states drive (SSD, 4x Samsung 860 EVO MZ-76E2T0B) for operating system and data storage.

### Data analysis and visualization

The image post-processing and visualization of the recorded data is done in four steps. In the first step, effective field of view of each tiled light sheet is cropped for each time point by a scripted macro in Fiji environment [30]. For example, for 6 tiles light sheet imaging, the cropping should be repeated 6 times (**Supp. Mat. Fig. 3**). In the second step, the cropped volume images are aligned and stitched together with the BigStitcher plugin [31] in Fiji. Then after, in order to gain the resolution, the fused data is deconvolved with Lucy-Richardson method by a GPU based plugin that is called CLIJ [32]. The final data visualization and segmentation is done with Fiji and Imaris 9.5 together.

### *Drosophila* culture and sample preparation

*Drosophila melanogaster* stocks were raised on corn-meal based media following standard procedures. Embryos were collected on yeasted apple juice plates between 2-3 hours after egg-laying. The embryos were dechorionated with 50 % commercial bleach for around 2 minutes followed by careful rinsing in tap water. Several embryos were selected and submersed into 1.5 % low melting agarose in Phosphate buffered saline (PBS) and rapidly transferred into a glass capillary with an inner diameter of 800 µm. The capillary was mounted to a holder and inserted into the imaging chamber. The temperature during imaging was 22°C. The following stocks were obtained from the Bloomington Drosophila stock centre (Bloomington, IN, U.S.A.): *y*^*1*^, *w*^*1118*^, *cv*^*1*^, *sqh*^*AX3*^; *P{w[+mC]=sqh-GFP*.*RLC}2* and *w[*]; P{w[+mC]=His2Av-EGFP*.*C}2/SM6a*. The *His2Av*.*EGFP* marker was introduced into a *klar*^*1*^ mutant background, which reduced diffraction of the illumination beam. The *ru*^*1*^,*klar*^*1*^ stock was a gift of M. Welte (Univ. Rochester, U.S.A.).

## Acknowledgements

We thank Ehsan Ahadi Akhlaghi, Jens Januschke, Peter Lehmann and Michael Welte for discussions throughout the work and critically reading the manuscript and their insightful comments. We thank CINSaT (Centre of Interdisciplinary Nanosstructure Science and Technology, University of Kassel) for support and discussion and would like to thanks Cristian Sarpe’s support of electronic and optical measurement equipment. We thank the *Drosophila* Stock Centre (Bloomington, U.S.A) for *Drosophila* stocks. We thank Birgitt Simon and Monika Winneknecht for preparing fly food and the workshop of the Natural Science Department of the University Kassel for constructing customized devices and tools.

## Competing interest

The authors declare that they have no competing interests. The University of Kassel (employer) has filed a patent application to the German Patent and Trade Mark office under file DE 10 2021 107 821.0 on the invention of the MT-SPIM microscope design and setup.

## Funding

This research was funded by startup funding from the University of Kassel to HAJM.

## Data availability

Primary imaging data can be made available upon request.

## Notes

### Competing Interest Statement

The authors have declared no competing interest.

